# In-Cell Structural Dynamics of an EGF Receptor during Ligand-induced Dimer-Oligomer Transition

**DOI:** 10.1101/458265

**Authors:** N. Kozer, A. H.A. Clayton

## Abstract

The epidermal growth factor receptor (EGFR) is a membrane protein that regulates cell proliferation, differentiation and survival, and is a drug target for cancer therapy. Ligand-induced activation of the EGFR kinase is generally regarded to require ligand-bound-dimers, while phosphorylation and downstream signalling is modulated by higher-order oligomers. Recent work has unveiled changes in EGFR dynamics from ligand-induced dimerization in membranes extracted from cells, however less is known about the changes in EGFR dynamics that accompany the ligand-induced dimer to tetramer transition in a live cell environment. In the present report, we determine the dynamics of a c-terminal GFP tag attached to EGFR in the unliganded dimer and in the liganded tetramer by means of dynamic depolarization microscopy. We made use of a novel analysis method, the single-frequency polarized phasor ellipse approach, to extract two correlation times on the subnanosecond and super-nanosecond timescales, respectively. EGF binding to the EGFR-GFP dimer lengthened the sub-nanosecond correlation time (from 0.1ns to 1.3ns), and shortened the super-nanosecond correlation time (from 210ns to 56ns) of the c-terminal GFP probe. The sub-nanosecond depolarization processes were assigned to electronic energy migration between proximal GFPs in the EGFR dimer or oligomer, while the super-nanosecond correlation times were assigned to nanosecond fluctuations of the GFP probe in the EGFR complex. Accordingly, these results show that ligand binding to the extracellular domain increased the average separation between the c-terminal tags and increased their rotational mobility. We propose that the dynamics are linked to an inhibitory function of the c-terminal tail in the un-liganded dimer and to the requirement of facile stochastic switching between kinase activation and cytoplasmic adaptor/effector binding in the active tetramer.

## 1. Introduction

The epidermal growth factor receptor (EGFR) is a membrane protein that regulates cell proliferation, differentiation and survival [1]. Overexpression or mutation of the EGFR occurs in 30% or more of cancers including brain, head-neck, breast, and lung cancer [2]. Consequently, the EGFR is considered an important target for rational drug therapy and there are a number of drugs that are used clinically to target the EGFR [3].

The EGFR is a single pass transmembrane protein and consists of an extracellular domain (ECD), a transmembrane domain (TMD), a juxta-membrane domain (JMD), a kinase domain (KD) and a cytoplasmic tail domain (CT). Activation of the EGFR is generally regarded to involve a ligand-induced oligomerization (dimerization) involving either a monomer-dimer transition [4], conformational change in pre-formed dimers [5] or higher-order oligomeric transitions [6–11]. Recent work has shown that the ligand-induced higher-order oligomers (tetramers and above) are phosphorylated [8,10,11] bind cytoplasmic adaptors [9] and modulate cell signalling [10,11]. Crucially, phosphorylated dimers appear to activate some signals but not others, indicating that either the structure and/or dynamics of the EGF-driven EGFR oligomers are responsible [10,11].

Structural studies using x-ray crystallography, NMR and biophysical techniques have produced models of the isolated fragments of the EGFR including the ECD [12–14], TMD [15,16], JMD [17,18], KD [19,20] and CT[21–23]. These models have provided static pictures of unliganded or liganded states of the ECD (i.e. tethered ECD monomer and liganded untethered ECD dimer), inactive and active states of the KD (symmetric inactive kinase dimer and asymmetric active kinase dimer), as well as, different conformational states of the TMD (N-terminal interface dimer versus C-terminal interface dimer) and the JMD. The CT domains appear to be intrinsically disordered [23].

While structural studies have revolutionized our understanding of EGFR conformations in the frozen state, measurements of EGFR conformational dynamics have the potential to provide significant insight into molecular recognition and allosteric transitions. Spectroscopic methods such as fluorescence, NMR, solid-state NMR, ESR/EPR and phosphorescence have provided valuable information on EGFR dynamics in solution, in synthetic membranes and in membranes extracted from cells. For example, fluorescence studies on isolated ECD and intracellular domain fragments (KD and CT) in solution provided evidence that ligand binding reduces the nanosecond internal motions in the ECD domain [24], while phosphorylation of the CT domain increases the nanosecond internal motions of the CT domain [22]. These studies are relevant to the question of whether ligand binding (or phosphorylation) induces a new conformation or selects a conformation from a preexisting conformational equilibrium. NMR studies on TM domains in micelles or synthetic membranes revealed rotational procession of the TM domains on the microsecond time scale, and nanosecond orientational fluctuations, consistent with the increased viscosity of the membrane [25]. Solid-state NMR [26] on the full-length receptor in membranes extracted from cells revealed that EGF binding decreased the dynamics of the ECD domain, while keeping the KD domain relatively rigid on the NMR timescale [26]. These results imply a conformational selection process for the ligand binding to the ECD domain as opposed to an induced fit mechanism. Furthermore, the loss in ECD entropy from the ligand binding was proposed to contribute to the energetics of the monomer to dimer transition. ESR [27] and time-resolved phosphorescence [28] studies probed microsecond to millisecond motions, which were assigned to uniaxial rotational procession of EGF-bound complexes. Collectively, these studies reveal rich dynamics that cover internal structural fluctuations, domain motions through to rotational procession of EGFR complexes on membranes.

To understand the structure and dynamics of the EGFR in a living cell requires biophysical measurements of the ensuing structures and dynamics in living cells. Somewhat surprisingly, there is a paucity of experimental data on how *EGF changes EGFR structural dynamics in a live cell environment (i.e. before and after EGF addition)*. In particular, nothing appears to be known about dynamics in the unliganded dimer and how ligand-induced dimer to tetramer (or higher-order oligomer) transition impacts on receptor structural dynamics.

In this paper, we address the above questions focussing on the CT domain of the receptor since it plays crucial roles in kinase inhibition and in coupling receptor activation to intracellular signalling cascades [21,22,42]. Specifically, we ask how dynamic is the CT domain in full length unliganded EGFR dimer in living cells? How does ligand-induced dimer to tetramer transition influence the dynamics of the CT domains in living cells?

To address these questions requires the use a cell system that contains high levels of pre-dimerized EGFR and liganded EGFR tetramers, and a microscopy method appropriate to the examination of structural fluctuations on the picosecond to sub-microsecond timescales on single living cells.

The BaF/3 system with stably transfected EGFR is a suitable system to examine the dynamics of the EGFR in pre-dimerized and ligand-bound tetrameric states. In our previous work, we showed the EGFR with a C-terminal GFP tag (EGFR-GFP) in BaF/3 cells was significantly pre-dimerized (greater than 90% EGFR dimer) [6,30] and after ligand binding was predominantly tetrameric [6]. Moreover, it was shown that the presence of the GFP tag does not interfere with the biological activity (phosphorylation [6], signalling and trafficking [31]) of the receptor. GFP-protein fusions appear in other studies of membrane protein structure and dynamics [32].

To measure the structural dynamics sensed by the fluorescent probes, we will use a new analytical method recently developed by us to extract complex fluorescence depolarization dynamics from fluorescently-tagged proteins in cells [33]. This method extends the earlier work of Weber [34], Gratton [35], Lakowicz [36] and Clayton [37] and utilizes steady-state anisotropy together with dynamic determination of depolarization using the frequency-domain technique. We have recently demonstrated that this method can resolve complex anisotropy decays and extract rotational correlation times in the range of sub-nanoseconds to hundreds of nanoseconds [33]. This time range is appropriate to examine structural fluctuations of membrane proteins without the interference from other depolarizing processes, such as uniaxial rotation about the membrane normal, which occur on the longer microsecond time-range.

The present paper is organized as follows. In the first section, we review our approach to the analysis of complex anisotropy decays, called the polarized phasor ellipse approach. In the second section, we then apply this analysis to steady-state and dynamic depolarization data obtained for EGFR-GFP [6,30] in BaF/3 cells in the absence and presence of ligand. In the third section, we discuss our results in the context of our current understanding of EGFR structure and dynamics.

## 2. Theory of the polarized phasor ellipse approach

Time-resolved anisotropy, whether measured in the time-domain or frequency-domain, is an established technique for measuring the hydrodynamics of biological macromolecules in solution [38]. This method measures a time-dependent change in the orientation of a fluorescent probe after photo-selection. The change in orientational distribution of photo-selected excited-states from an ensemble of macromolecules is recorded using anisotropy decay (time-domain) or dynamic depolarization (frequency-domain).

The decay of the anisotropy with time is usually analysed as an exponential (or sum of exponentials). In this case, the anisotropy decays (r(t)) from an initial value at zero time (r(0)=r_0_) to some other value over time (equation 1). For the simplest anisotropy decay, r(t) is given as a single exponential function

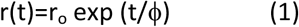

where ϕ is the rotational correlation time.

We utilise the dynamic depolarization approach with wide-field imaging [39]. This approach allows anisotropy decay information to be extracted from many cells in parallel and with good photon efficiency. Until recently, dynamic depolarization data recorded with a single modulation frequency could only be analysed in terms of very simple anisotropy decay models (single exponential anisotropy decay or single exponential anisotropy decay plus an offset anisotropy) [33, 34, 39]. However, using a new approach, called the polarized phasor approach (also called polarized AB plot [37]), and with certain restrictive conditions (known lifetime and known limiting initial anisotropy), we demonstrated that we could extract two correlation times in the range of sub-nanosecond and tens of nanoseconds from a fluorescent probe in living cells [33]. The full theory of this approach appeared in an earlier publication [33]. Here we provide a brief summary only.

The phasor is a vector that represents phase and modulation on a two-dimensional plot. When dealing with dynamic depolarization data recorded in the frequency-domain, the phasor from the perpendicular-polarized component is used. The result from a single frequency experiment, then, is one point on a 2D plot. To interpret the anisotropy decay we also draw something called the polarized-phasor ellipse. This ellipse traces out the trajectory (in phasor space) corresponding to all single exponential decaying anisotropies for the system under consideration (in the time-domain this means equation 1, fixed r_0_ and variable ϕ). With the polarized phasor approach, the complexity of the anisotropy decay (simple, one correlation time; complex, more than one correlation time) can be inferred by inspection of the position of the experimental data relative to the polarized phasor ellipse (on the ellipse, single correlation time, inside the ellipse, more than one correlation time). Moreover, the graphical approach enables multiple correlation time models to be assessed using geometry and vector algebra. For example, the sum of two exponentially decaying anisotropies (i.e. two correlation times) is represented on the graph by a line that intersects with two points on the ellipse and with the experimental data point. The points of intersection with the line represent the two correlation times and the relative distance of the experimental point to each intersection to the ellipse is related to the fractional contribution of each correlation time to the anisotropy decay.

The polarized phasor ellipse is plotted as an x-y graph, with natural coordinates for this type of system, G=mcosp and S=msinp where m is the modulation and p is the phase. Given a fixed limiting zero-time anisotropy of r_0_, a fixed excited state lifetime of τ_2_ and a fixed modulation frequency of ω, the phasor components from perpendicular-polarized emission signals are given by,

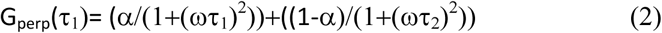

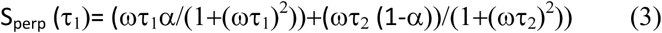

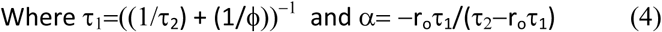

An example of a perpendicular polarized phasor ellipse plot is shown in Figure 1 for r_0_=0.4, τ_2_=4ns, ω=40MHz over the rotational correlation time range (ϕ=0.1,0.2,0.5,1,2,5,10,20,50,100 ns). The G_perp_ versus S_perp_ plot traces out an elliptical shape referred to as a lemniscate (a ribbon). A simulated double exponential anisotropy decay with correlation times of 0.5 ns and 20 ns and fractional fluorescence contribution of the 0.5ns component, β, is in Figure 1. It is clear that the point corresponding to the double exponential anisotropy decay is located inside the ellipse, as expected. Drawing a line between the polarized phasor positions corresponding to 0.5 ns and 20 ns reveals that the simulated data point lies on the line and closer to the phasor corresponding to the 0.5ns, as predicted from the simulation.

**Figure 1.**
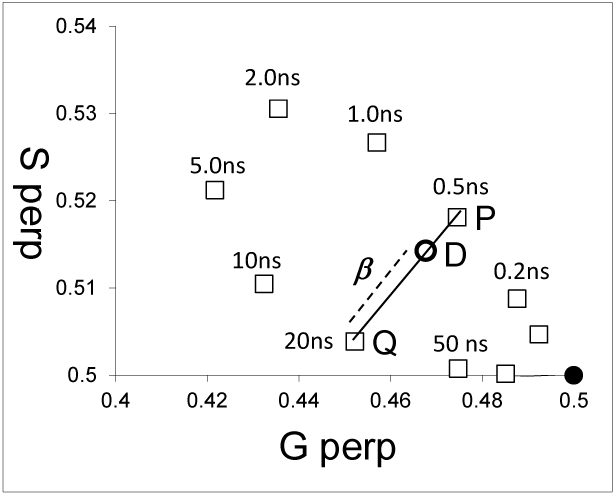
Polarized phasor ellipse plot. Open squares reveal positions of the perpendicular-polarized phasor computed for single correlation time rotors (r_0_=0.4 and τ_2_=4ns, ϕ varies from 0.01 ns to 100ns).

From the two correlation times, ϕ_1_ and ϕ_2_ and the fractional fluorescence β (r_0_ and τ_2_ fixed), we can compute the steady-state anisotropy, r, viz,

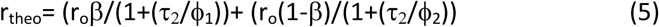

The way we implement the polarized phasor approach in practice is as follows. First, given r_0_ and τ, we construct the polarized phasor ellipse using equations 2–4. Second, measurements of single-frequency dynamic depolarization yield the cosine and sine components of the perpendicular-polarized phasor. We then plot this data point, D (G_perp_,S_perp_), on the polarized phasor ellipse. If it lies on the ellipse, we can deduce that the depolarization process is simple (one correlation time) and the value of the correlation time deduced from the location of experimental point on that ellipse. If it lies inside the polarized phasor ellipse then the anisotropy decay is complex and a double exponential decay for the anisotropy is considered. The double exponential anisotropy decay is analysed using the constraints provided by the polarized phasor ellipse and vector algebra. First, a value for ϕ_1_ is guessed and using equations (2)–(4), the theoretical phasor position P(G_perp_ (ϕ_1_),S_perp_ (ϕ_1_)) corresponding to ϕ_1_, is computed. The second correlation time (ϕ_2_) is calculated from the corresponding phasor Q(G_perp_ (ϕ_2_),S_perp_ (ϕ_2_)) from the condition of collinearity between the points D, P and Q. This is achieved computationally by varying the value of ϕ_2_ so as to minimize the difference between the gradients of the lines DP and QP. From the ratio of lengths of two line segments (length (DP)/length (PQ)), we can deduce β. Then we can calculate the theoretical steady-state anisotropy corresponding to ϕ_1_, ϕ_2_ and β from equation 5. The theoretical steady-state anisotropy is compared with the experimental steady state anisotropy and a goodness-of-fit, computed. The goodness-of-fit is given by the equation,

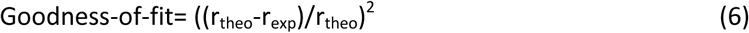

A different value for ϕ_1_ is then chosen, a different set (ϕ_2_, β) extracted, thence a new r_theo_, and goodness-of-fit computed. This iterative approach repeats until the parameters with the lowest goodness-of-fit are found. It is important to note that in this minimization, the only independent variable is one of the correlation times (either ϕ_1_ or ϕ_2_). The other correlation time and the fractional contribution determined by the constraints of geometry imposed by the polarized phasor ellipse and the experimental data point. Moreover our “one parameter fit” approach differs substantially from the conventional analysis of anisotropy decay data where typically 3-4 parameters are independently varied (r_0_, β, ϕ_1_, ϕ_2_).

## 3. Materials and Methods

### 3.1 Materials

The murine hemopoietic cell line BaF/3 expressing C-terminally tagged EGFR-GFP constructs (Ludwig Institute for Cancer Research, Melbourne, Australia) has been described previously [6,8,9,30]. Experiments were carried without and with epidermal growth factor in living cells under conditions where the EGFR-GFP was kept at the cell surface, as described in ref 9 to avoid complications associated with receptor internalization and trafficking.

### 3.2 Methods

#### Data Analysis

The dynamic depolarization microscope and associated methods (including corrections for G-factor, aperture depolarization and background) has been previously described [33]. Data collected from the dynamic depolarization microscope were (i) steady-state anisotropies from individual cells, (ii) excited-state lifetimes from individual cells and (iii) perpendicular-polarized phasors from individual cells. These data were averaged to obtain cell-population-averaged lifetime, average steady-state anisotropy and averaged perpendicular-polarized phasors. Data analysis consisted of assuming a particular model for the anisotropy decay, representing that model on the polarized phasor plot, computing the steady-state anisotropy associated with that model and then comparing the theoretical steady-state anisotropy with the experimentally-measured steady-state anisotropy. The three models considered were isotropic rotator, hindered rotator, and double exponential anisotropy decay. Triple exponential anisotropy decay analysis and Monte Carlo simulations of energy migration are described in the Appendix or Supplementary Materials. Some of the steady-state anisotropy data from EGFR-GFP in the absence of ligand has already been published [30] and listed here in Table 1.

**Table 1.**
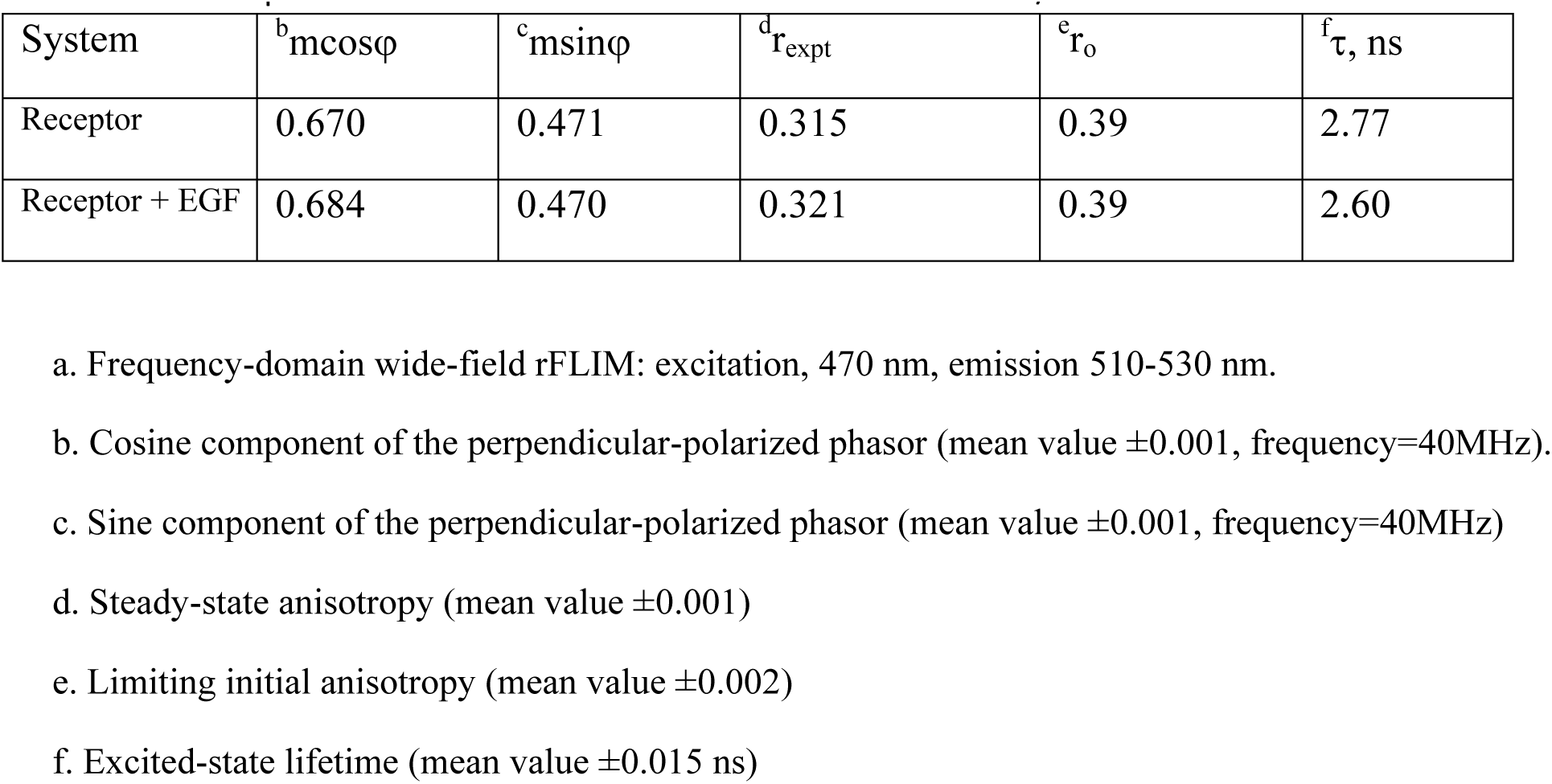
rFLIM^a^ parameters for EGFR-GFP and EGF-EGFR-GFP in BaF/3 cells.

## 4. Results

Using our wide-field frequency-domain dynamic depolarization microscope, we measured lifetime, steady-state anisotropy and perpendicular-polarized phasor components of GFP fluorescence from stably-transfected EGFR-GFP construct in BaF/3 cells. Some of the steady-state anisotropy data from EGFR-GFP in the absence of ligand has already been published [30] and reproduced here. Pertinent results are in Table 1.

In the absence of EGF, the EGFR-GFP emission from BaF/3 cells was depolarized to some extent (r_expt_<r_0_). Inspection of the experimental perpendicular polarized phasor relative to the polarized phasor ellipse plot, revealed that the phasor was inside the polarized phasor ellipse (Figure 2A). Assuming some systematic error, the closest single correlation time to the data was 200ns. The theoretical steady-state anisotropy (equation 5 with r_0_=0.39, β=l, ϕ_1_=200ns, τ=2.77ns Table 2) for this simple model was 0.385, which does not agree with the measured steady-state anisotropy of 0.315. A one sample t-test revealed that the difference between theory and experiment was extremely statistically significant (Table 1, p<0.0001, mean (expt)=0.315, s.d. (expt)=0.017, N=307). Moreover, the location of the experimental polarized phasor inside the polarized phasor ellipse plot, strongly indicated a more complex mode of depolarization. Therefore, we can reject a single correlation time model for the anisotropy decay from EGFR-GFP in cells. We next consider a hindered-rotator anisotropy decay model.

**Figure 2.**
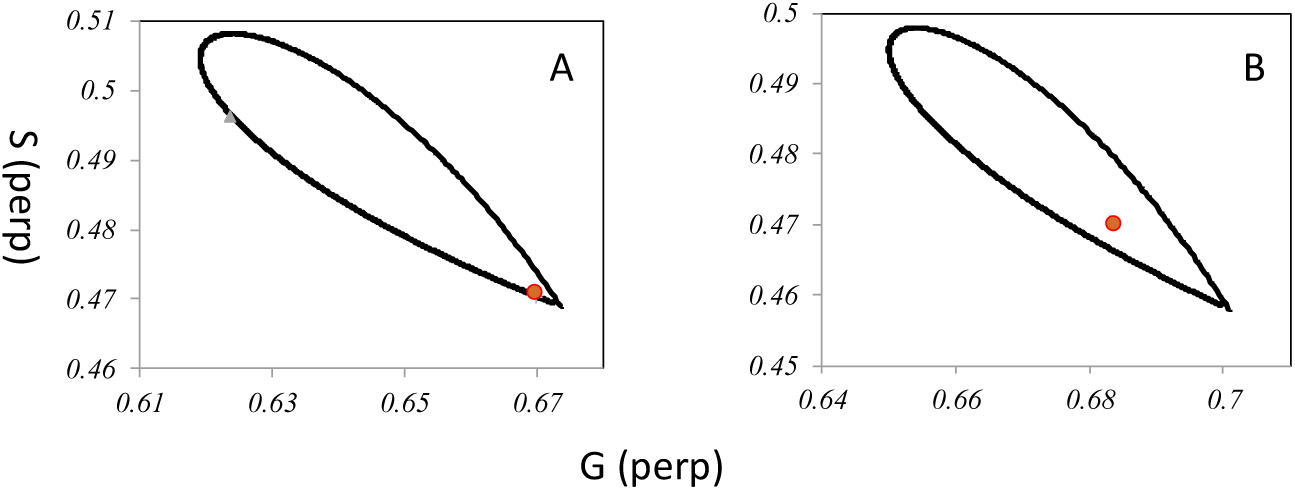
A. Polarized phasor ellipse plot for EGFR-GFP (r_0_=0.39 and τ_2_=3.77ns, ϕ varies from 0.01 ns to 100ns) and experimental data point for perpendicular-polarized phasor from EGFR-GFP in BaF/3 cells. B. Polarized phasor ellipse plot for EGF-EGFR-GFP (r_0_=0.39 and τ_2_=3.6ns, ϕ varies from 0.01 ns to 100ns) and experimental data point for perpendicular-polarized phasor from EGF-EGFR-GFP in BaF/3 cells.

**Table 2.** Time-resolved fluorescence anisotropy decay models and extracted parameters.

| Construct | Model | $\phi_1$, ns | $\phi_2$, ns | $\alpha_1$ | $\alpha_2$ | $r_\infty$ | $r_{\text{theo}}$ | $r_{\text{expt}}$ |
| --- | --- | --- | --- | --- | --- | --- | --- | --- |
| EGFR-GFP | Isotropic | 200 | - | 1 | 0 | 0 | 0.385 | 0.315 |
| EGFR-GFP | Hindered | 6.8 | - | - | - | 0.362 | 0.382 | 0.315 |
| EGFR-GFP | Double | 0.07 | 209 | 0.185 | 0.814 | - | 0.315 | 0.315 |
| EGFR-GFP<br>+EGF | Double | 1.29 | 55.80 | 0.21 | 0.79 | - | 0.321 | 0.321 |

The simplest double exponential anisotropy decay model is the hindered rotator, with the second rotational correlation time being essentially infinite (ϕ_2_=∞). Drawing a line through the experimental phasor and the polarized phasor ellipse position corresponding to (ϕ_2_=∞) we obtained ϕ_1_=6.8ns, and β=0.7 from intersection of the line with the polarized phasor ellipse. Using equation 5 and these parameters (r_0_=0.39, ϕ_1_=6.8ns, ϕ_2_=∞,β=0.7) we computed a theoretical steady-state anisotropy of 0.382. A one sample t-test revealed that this theoretical value for the steady-state anisotropy was significantly different from the experimental value (Table 1, p<0.0001, mean (expt)=0.315, s.d. (expt)=0.017, N=307). Therefore, we can reject the simple hindered rotator model for the depolarization process from EGFR-GFP on BaF/3 cells.

The next level of complexity is the double-exponential anisotropy decay model allowing both correlation times to be finite. Using our approach, we scanned possible values of ϕ_2_ and extracted (ϕ_1_,β). We then computed r_expt_ and goodness-of-fit parameter using equations 5 and 6. A plot of the goodness-of-fit parameter as a function of ϕ_2_ and ϕ_1_ is in Figure 3. The plot reveals a single minimum corresponding to the best fit. The model with the lowest goodness-of-fit parameter has the parameters (ϕ_1_=0.1ns, ϕ_2_=210ns, β=0.l8, Table 2). In the context of a double exponential anisotropy decay model we extracted two correlation times corresponding to sub-nanosecond and super-nanosecond depolarization processes. As expected from the requirements of the model minimization process, the computed theoretical anisotropy corresponding to this model was 0.315, in excellent agreement (i.e. better than 1%) with the experimental anisotropy of 0.315. A one sample t-test revealed that we could not reject the null hypothesis that the theoretical anisotropy and experimental anisotropy are the same (Table 1, p=1, mean (expt)=0.315, s.d. (expt)=0.017, N=307).

**Figure 3.**
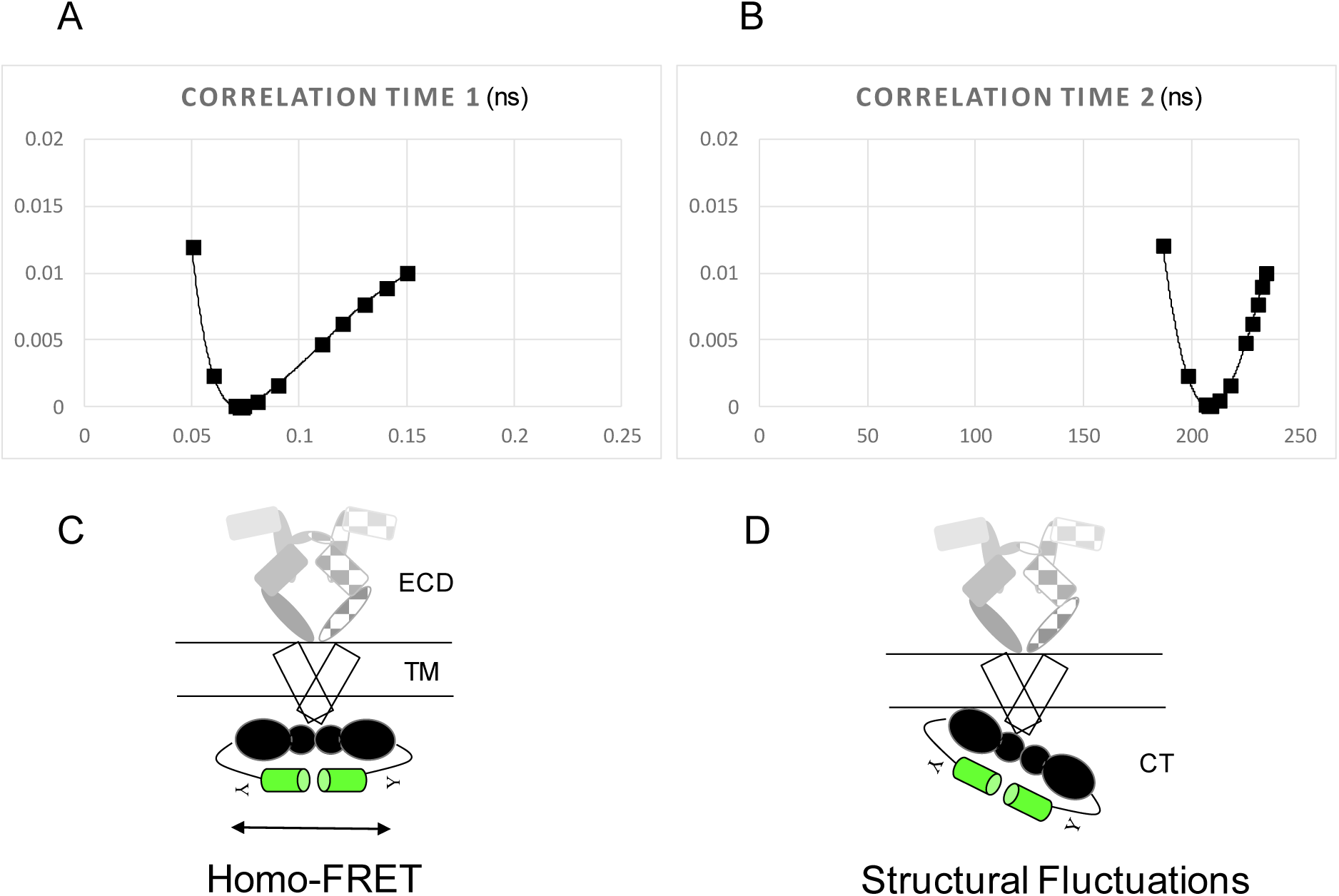
Double-correlation time model for EGFR-GFP dimer in BaF/3 cells. A. Chi-squared plot for correlation time 1. Note the single minimum at 0.1ns B. Chi-squared surface for correlation time 2. Note the single minimum at 209ns. C. Tentative model for the unliganded EGFR dimer. Arrows indicate energy migration between GFP tags. D. Tentative model for the unliganded EGFR dimer. Arrows indicate motions of the GFP tags.

Addition of EGF to EGFR-GFP in the BaF/3 cells led to a change in depolarization processes as reflected in the parameters collected in Table 1 and in the position of the perpendicular polarized phasor with respect to the polarized phasor ellipse (Figure 2B). The steady-state anisotropy and cosine component of the perpendicular-polarized phasor increased, while the excited-state lifetime decreased slightly (but remained in the range of lifetimes reported for GFP-containing proteins in cells). In the context of a double exponential anisotropy decay model, as for EGFR-GFP, the best-fit parameters for the liganded complex were (ϕ=1.3ns, ϕ_2_=56ns, β=0.21). The goodness-of-fit surfaces for this model are shown in Figure 4. The steady-state anisotropy from the theoretical model was 0.321, in excellent agreement with the observed experimental anisotropy of 0.321 (Table 1, p=1, mean (expt)=0.321, s.d. (expt)=0.025, N=119) .

**Figure 4.**
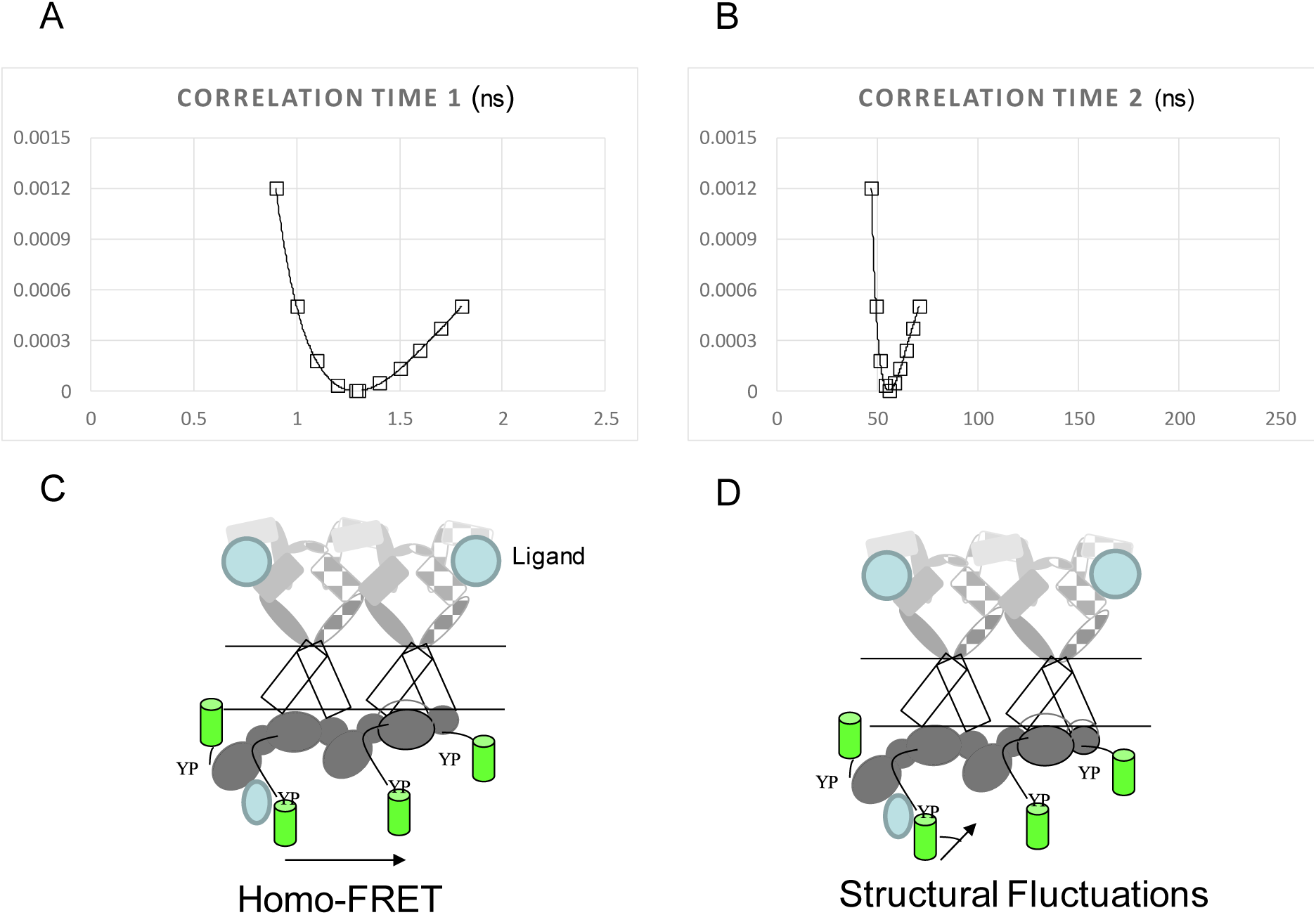
Double-correlation time model for EGF-EGFR-GFP tetramer in BaF/3 cells. A. Chi-squared plot for correlation time 1. Note the single minimum at 1.3ns B. Chi-squared surface for correlation time 2. Note the single minimum at 56ns. C. Tentative model for the liganded EGFR tetramer. D. Tentative model for the liganded EGFR tetramer.

To summarize, EGF addition to EGFR-GFP to form the EGF-EGFR-GFP complex increased ϕ_1_ from about 0.1 ns to 1 ns and decreased ϕ_2_ from about 200 ns to 50 ns. In the following discussion section, we assign these correlation times and discuss their significance in the context of EGFR structure and dynamics.

## 5. Discussion

### Assignment of depolarization processes

We begin by discussing the short correlation times for the EGFR-GFP and for the EGF EGFR-GFP complex. The short depolarization times observed for the GFP probe attached to the EGFR in EGFR-GFP (ϕ_1_=0.1ns) and in the EGF-EGFR-GFP complex (ϕ_1_=1ns) are too short to be due to dynamic rotation of the GFP probe. The rotational correlation time of free GFP is about 10-20 ns in the cytoplasm of bacterial cells [39], which is at least an order of magnitude longer than the short depolarization correlation times observed here. The most likely explanation is that the short depolarization process is due to energy migration (homo-FRET) between proximal GFPs. Evidence for homo-FRET was previously reported for EGFR-GFP on BaF/3 cells, through an enhancement of steady-state anisotropy during progressive photobleaching [30]. The fraction of steady-state anisotropy lost due to homo-transfer (β=0.16) estimated from that study [30] agrees well (i.e. to within 15%) with the fractional anisotropy due to the short correlation time from the present dynamic depolarization measurements (β=0.18, Table 1). Gerritsen and colleagues also interpreted rapid depolarization (sub-nanosecond) due to homo-FRET using a time-gate polarization microscopy method for EGFR-GFP on NIH 3T3 cells [40].

The long correlation times observed for EGFR-GFP and the EGF-EGFR-GFP complex are many fold larger than the rotational correlation time for free rotation of GFP. Therefore unrestricted local probe rotation about a free tether cannot account for the magnitude of the correlation times. On the other hand, free uniaxial rotation of membrane proteins about the membrane normal have correlation times on the microsecond timescale [27,28]. Therefore, the long rotational correlation times measured here are likely to be due to internal nanosecond structural fluctuations of the EGFR (or possibly semi-independent motions of EGFR sub-domains) sensed by the GFP probe.

### Comparison with structural models for EGFR inactive kinase dimer

There is presently no high-resolution structure for the full-length inactive EGFR dimer in a living cell membrane. Structural data on the KD-CT fragment indicate a symmetric kinase dimer with an incompletely-resolved CT tail [18,41]. Recent biophysical studies on the isolated CT tail in solution point to a lack of significant secondary structure with features characteristic of an intrinsically disordered domain [23]. If we assume (initially) no significant interaction of the CT domain with the KD domain then we would expect the GFP probe to sample a large amount of space at the cytoplasmic side of the membrane. To compare this notion with experiment, we made a model where the two CT tails in the EGFR dimer are unstructured, free, and emanate from the ends of a symmetric kinase dimer. We generated a Monte Carlo simulation of probe-pairs allowing for random probe orientation, random probe distance from the KD (up to the maximum length of CT domain of 15nm from the Kratky plot in reference 23) and a fixed separation from the points of attachment to the KD domains of 5nm or 10nm (see Appendix or Supplementary Results). The results of this simulation are shown in Table S3 with the key comparison between the magnitudes of the theoretical steady-state anisotropy. It is note-worthy that the assumption of free chains cannot reproduce the experimental energy migration results for the EGFR-GFP dimer. The results imply that the CT tails are somewhat restricted in the EGFR-GFP dimer perhaps because of interactions of the CT tail with the KD domain of the receptor (or some other component on the cytoplasmic side of the membrane).

In the crystal structure of Jura et al [18] there are contacts between parts of the CT domain and the kinase dimer interface. Residues 967 to 978 form an alpha helix packed against the neighbouring kinase while residues 979-981 form a salt bridge with residues on the neighbouring kinase. Although the exact position of the C-terminus is not resolved in that structure, FRET studies from Koland's lab between a C-terminal BFP (a GFP variant, blue fluorescent protein) and an acceptor-labelled ATP revealed close approach between the c-terminal BFP of the CT domain and the ATP binding site on the kinase domain [22]. Studies of engineered CT tails with variable lengths led to the conclusion that the C-terminus loops back towards to the KD instead of extending away from it [22]. Given the distance between ATP binding sites is 3nm in the symmetric kinase dimer [41], we would expect a close proximity between GFPs in a EGFR-GFP dimer if the CT domain C-terminal GFP probe interacts substantially with the KD near the ATP binding site.

From our dynamic depolarization experiments, we can gain estimates of the distances between GFPs in the EGFR-GFP dimer in the context of a single distance model. From the correlation time of ϕ_1_=0.1ns, the energy migration rate in the EGFR-GFP dimer, k_em_, is given as k_em_=(1/2ϕ_1_)=5×10^9^ s^−1^. The inter-GFP distance, R, is then given by R=(1/ k_em_τ_2_)^(1/6)^ R_0_, which with R_0_=4.9±0.4 nm (orientation factor limits for fixed dipoles) and τ_2_=3.8×10^−9^ s (Table 1), we obtain R=0.61×R_0_=3.0±0.2nm. This is evidence, then, that the two CT tails in the EGFR-GFP dimer are located at close proximity to each other as predicted from the considerations above. A speculative model for the dimer is shown in Figure 3 assuming a untethered back-to-back ectodomain dimer and a symmetric head-to-head kinase dimer. Note the positions of the GFP tags close to the ATP binding sites.

The looping-back model was also tested using Monte Carlo simulations allowing for two-probes to be located on the surface of the kinase dimer (see Appendix/Supporting Material). This model gave good agreement with experiment provided the probes were constrained to the N-terminal halves of the kinase domain, closest to the kinase-kinase interface.

Evidence of interactions in the EGFR-GFP dimer can also be gauged by examination of the magnitude of the rotational correlation time associated with GFP motion. As noted above, the measured correlation time is a factor of at least 10 greater than the rotational correlation time of free GFP (ϕ_2_=210ns >>ϕ_GFP-free_=20ns). Since the GFP is attached via the CT domain to the EGFR, interactions between the CT domain and the KD domain could lead to reduced rotational diffusion and increased rotational correlation time. A rough estimate of the size of the rotational unit can be gleaned with an equivalent sphere approximation. Using ϕ=0.6 ns/kD and ϕ_2_=210ns, we obtain an estimated effective molecular weight of 210ns kD/0.6 ns=350 kD for free rotation. The apparent molecular weight derived from the published Stokes radius of the isolated unphosphorylated KD-CT domain dimer is 313±17 kD [42], which appears to be fortuitously close (i.e. <15%) to the value estimated from our dynamic depolarization data (adding two GFPs gives a MW of 313+54=367 kD). The implication, from this admittedly crude analysis, is that the two KD-CT domains in the EGFR dimer rotate or undergo orientation fluctuations as a single entity, which may occur if the JMD (in the inactive EGFR dimer) provides a loose linkage between the TM domain and the KD dimer. There is some evidence that this may occur in electron microscopy images of nearly-full length dimeric EGFR in membranes where symmetric and asymmetric KD dimer domains exist in multiple orientations [43]. Alternatively, the 200ns correlation time may represent other internal motions sensed by the GFP probe in the EGFR-GFP. For example, interactions of some or all the intracellular domain (i.e. juxtamembrane domain) with membrane components in the inner leaflet of the cell membrane could restrict the KD-CT domains and account for the lengthening in the GFP correlation time (without formation of any specific kinase dimer). A precise assignment awaits further experimentation and molecular dynamics simulations of the complete molecule in a membrane environment.

### Comparison with structural models for EGF-EGFR tetramer

As for the EGFR dimer, there is currently no structure of the full length EGFR activated dimer or tetramer in living cells. However, crystal structures of the kinase domain indicate an asymmetric dimer as the kinase-activated species [20]. In this structural model, there is a substantial reorientation of the inactive kinase dimer from a head-to-head orientation to a head-to-tail orientation [18, 20]. Models built from microscopy studies suggest that the asymmetric kinase dimers can stack as cyclic tetramers [6, 7, 8], extended tetramers, hexamers and higher-order oligomers [10,11].

Biophysical studies on isolated KD-CT domains in solution indicate that tail phosphorylation induces a conformational change in the CT domain increasing its local mobility and partially releasing it from the KD [21, 22].

The observed increase in short correlation time for the EGF-EGFR-GFP complex from 0.07ns to 1.2ns indicates that the efficiency of energy migration (homo-FRET) between GFP tags has decreased in the ligand-receptor complex relative to the un-liganded receptor complex in the cellular environment.

Proceeding along similar lines to that of the EGFR-GFP unliganded dimer, we ran a Monte Carlo simulation assuming that the CT domains in the liganded tetramer released from the KD domains, and essentially free. This model predicted an anisotropy which had a discrepancy of 15% with the experimental value. The looping-up model was also tested and produced a larger discrepancy with the experimental data of 30%.

An alternative model posits that the GFPs are held in a single defined geometry. If we assume energy migration only occurred between only two GFPs in the EGF-EGFR-GFP complex and that the Forster distance was unchanged in the unliganded and liganded complexes, the distance between GFPs would increase by a factor of (1.2/0.07)^^1/6^=1.6, which increases the separation between GFP tags from R= 3nm to R= 4.8nm.

However, consideration of the energy migration formalism between 4 dipoles (4 GFPs) in a tetramer leads to the following equation for the time-dependent anisotropy decay for a square planar geometry in the limit of random dipole orientations,

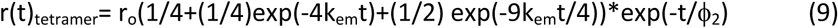

In this case, the observed short correlation time, ϕ_1_ would be a weighted average of the energy migration rates and ϕ_1_ could be between 1/4k_em_ and 4/9k_em_. The distance between two neighbouring GFPs in the tetramer calculated with these limits is in the range 5-6.5 nm. We conclude irrespective of the energy migration model, that there is an increase in the average separation of GFP tags during the ligand-induced dimer to tetramer transition. This suggests a change in the conformation of the receptor C-terminus during and/or after activation. This observation agrees well with the conclusions from the Pike lab. The Pike lab used luciferase complementation of probes fused to the c-terminal tails of the EGFR in CHO cells [44]. Evidence for an increase in the separation between the c-terminal tags was found following EGF stimulation and EGFR kinase activation by an initial decrease in luciferase signal [44]. The increase in separation could result from the change in kinase conformation from symmetric to asymmetric dimer, release of the CT tail from the kinase dimer or a combination of the two processes.

Insights into possible release of the CT tail from the KD domain can also be disclosed by consideration of the rotational correlation time of the GFP in the EGF-EGFR-GFP system. The magnitude of the rotational correlation of 50ns signifies a significant increase in rotational fluctuations or motion of the GFP tag in the activated EGFR-GFP tetramer relative to the inactive EGFR-GFP dimer (200ns). In studies of the isolated monomeric KD-CT fragment Koland's lab noted an decrease in rotational correlation time from 86 ns in the unphosphorylated state, to 56 ns in the phosphorylated state, which qualitatively agrees with the decrease in rotational correlation time here [22]. If we assume the CT tail is phosphorylated, largely unstructured and released from the KD region in the activated EGFR-GFP tetramer, the magnitude of the rotational correlation time reports on the MW of the species attached to the phosphorylated CT tail. If we make the same calculation of effective molecular weight as we did above, we obtain an apparent molecular weight of 50/0.6 KD =83 kD for the size of the freely rotating unit. This molecular weight is too small to be due to free motion of the KD-CT domain tetramer, dimer or monomer indicating increased local motion of the CT tail. However, the molecular weight is too large to be due to a completely free GFP (27kD) which means there is still some restriction to rotational motion of the CT tail sensed by the GFP. We suggest that the recruitment of adaptors [9] and other enzymes to the CT tail after activation has partially impeded the rotation of the GFP probe. A tentative model of the EGF-bound tetramer is shown in Figure 4 showing partially-released CT tails and a bound cytoplasmic protein. The arrangement of the extracellular domains is from a recently published model of the EGFR tetramer. We conclude that in the activated tetramer, the CT tails are, on average, less restricted in spatial organization or motion than in the inactive dimer.

### Alternative explanations

Our discussion above appears to be consistent with current knowledge of the EGFR structure and dynamics, particularly of analogous fluorescence studies on the isolated CT domain [21,22]. However, it is important to consider alternative interpretations of our data.

1. Homo-FRET is due to dimerization between GFP tags. One could envisage the short correlation times are due to energy migration between the GFP probes as part of a GFP-GFP dimer, instead of an EGFR-mediated dimer. The close distance between GFP probes measured here could potentially be due to GFP-GFP dimerization since the dimensions of a single GFP is cylindrical with a short axis of about 2nm and a long axis of 4nm. However, the calculated energy migration rate for such a dimer is k_em_=400×10^9^ s, which is a factor of 50 greater than the measured rate for energy migration in EGFR-GFP here. Moreover, attempts to the fit the data to a model with a ϕ_1_=2.5 ps yielded a goodness-of-fit parameter which was a factor of 10^8^ larger than the model with ϕ_1_=70 ps. On this basis, therefore, we believe we can exclude GFP-GFP dimerization as the sole source of the energy migration in this instance.
2. Rotational diffusion model for GFP motions is inappropriate. An alternative model due to Weber [45] posits that the anisotropy decay is not from rotational diffusion but represents a stochastic switching between alternative fixed orientations of the probe molecule. This situation has been further discussed in the dynamic depolarization literature [46]. However, this may be relevant to the case here where the CT tail of the EGFR is required associate with the kinase domain during inhibition (and during enzyme catalysis) and dissociates after phosphorylation where it can interact with cytoplasmic adaptors or effectors. If we model the movement as a binding/un-binding stochastic process between two orientations (say one in which the GFP is close to the KD domain and the other when the GFP is away from the KD domain), as in the model of Weber, the rotational correlation time is related to the Markovian switching time between the two conformational states. If k_12_ represents the rate of conformational change between orientations 1 and 2 and k_21_ represents the reverse rate, the rotational correlation time is given by ϕ_2_=1/(k_12_+k_21_). If k_12_>>k_21_, then k_12_=1/ϕ_2_=5×10^6^ s^−1^ and if k_12_=k_21_, then k_12_= k_21_=1/2ϕ_2_=2.5×10^6^ s^−1^. Thus, sub-microsecond scale stochastic switching of the CT tail (perhaps between free and KD-bound states) could alternatively account for the observed dynamics in the EGFR-GFP dimer. An analogous calculation in the EGF-EGFR-GFP tetramer reveals possible switching rates of k_12_=1-2×10^7^ s^−1^, which are an order of magnitude faster than in the EGFR-GFP dimer.
3. The GFP tag is too perturbative to probe motions of the CT tail of EGFR. The size of the GFP tag is 27kD, which is equivalent to a large domain in a protein. Therefore, it is conceivable that the tag is too large to reliably report on the dynamics of the CT tail of EGFR. Functional biochemical studies of EGFR-GFP in cells suggest, somewhat surprisingly, that the EGFR-GFP is still functional in terms of phosphorylation, coupling to down-stream effectors close to the C-terminal tail, and trafficking. Thus, functionally, the GFP is not perturbative when judged by biochemical methods. However, because of the size of the GFP, the fastest motional depolarization is 10-20 ns (the motion of free GFP) and thus we cannot provide information on sub-nanosecond rotational motions (or fluctuations) of the CT tail that might be important for functioning. (However, we note parenthetically, that of course this is the reason why GFP is so useful for measuring homo-FRET). We cannot quantitatively assess the degree to which the GFP tag perturbs the CT dynamics from our current measurements and this quantitation awaits further experiments with smaller probes and molecular dynamics simulations.
4. Simplicity of anisotropy decay model. The analytical approach we employed allows distinction between single correlation time, hindered rotator and double correlation time models (i.e. without recourse to multiple frequencies). With our current analysis of the EGFR-GFP we have a single correlation time due to energy migration and another correlation time due to rotation. It is unlikely that a single correlation time for either process can capture the complexity of the environment sensed by the probe in a complex cellular environment. We considered the possibility that a triple exponential correlation time model could account for the experimental data. Using a parameterized three-correlation time models (see appendix for details) with correlation times on the sub-nanosecond, nanosecond and super-nanosecond timescales we could get good fits to our experimental data. For the EGFR-GFP dataset, attempts to fit to three yielded very small amplitudes for the nanosecond magnitude correlation time. This suggests that for a sum of exponential anisotropy decays model, a double exponential seems to be the most resolvable model with our EGFR-GFP data and current approach. For the EGFR-GFP dataset we could fit to alternative models which included nanoseconds, tens of nanoseconds and hundreds-of-nanoseconds timescales. We interpret this model flexibility to indicate that the EGF-EGFR-GFP data could be described by a more complex model than a double exponential but a unique triple exponential model could not be resolved. We wish to stress that our data and analysis approach does not permit fitting to more complex models, which would require multi-frequency datasets (or high-resolution time-resolved fluorescence). However, the anisotropy function and the components of the polarized phasor have the property that they are linear in the fractional fluorescence contributions of different species. This property means that any distribution of states can be *simulated* for comparison with experiment (e.g. results from Monte Carlo or molecular dynamics simulations). For example, we showed that a random chains model could not account for the energy migration data from the unliganded EGFR-GFP dimer but a model of a looped-up CT tail provided a better description.

### Mechanistic insights

Recent biophysical studies on the isolated EGFR CT domain indicate that it possesses the properties of an intrinsically disordered protein [23]. In this context, ligand-induced modulation of CT dynamics in the full-length EGFR protein may be a means to influence biological functionality. Our results show here for the first time that CT dynamics is restricted in the unliganded dimer but less restricted in the liganded oligomer, while previous NMR [26] and fluorescence studies on the ECD domain [24] (isolated or as part of full length EGFR in membranes) indicated the opposite trend in ECD dynamics upon EGF binding. We propose, in line with Koland [21,22], that in the inactive EGFR dimer, the CT domain is in a predominantly auto-inhibited state via a "reversible latch" in contact with the KD domain. Ligand binding to the ECD transfers entropy from the initially flexible ECD domain to the relatively rigid CT domain. This increase in CT domain dynamics may help to release the CT domain, and allow more efficient cytoplasmic target search capabilities in the context of conformational selection and fly-casting mechanisms [23]. The importance of understanding the structure and dynamics of the EGFR tetramer (as opposed to the dimer) is underscored by recent studies showing the insufficiency of the phosphorylated EGFR dimer in activating Ras [47] and the importance of EGFR oligomers initiating signalling from the cell surface [9,10,11,48]. We propose that the inherent flexibility (and spatial arrangements) of the CT domains in the EGF-activated tetramer may allow efficient search, engagement and recruitment of adaptors/effectors to effectively engage and activate cytoplasmic partners.

## 6. Conclusion

We have shown that the nanometre spatial disposition and nanosecond motions of a GFP probe attached to the CT domain of full length EGFR are restricted in the EGFR pre-dimer and less restricted in the EGF-EGFR-GFP tetramer (or higher-order oligomers) in a live cell environment. We propose that these motions reflect the stable interactions between the CT domain and KD domain during inhibition in the EGFR-GFP dimer and transient interactions between the CT domain, KD domain and cytoplasmic adaptors required during enzyme catalysis and adaptor binding in the EGF-EGFR-GFP tetramer. The increase in the motions of the CT tail and the decrease in energy migration observed in the activated tetrameric complex are in accordance with a model in which the CT tail becomes displaced from its interaction with the KD and samples a greater conformational space.

## 7. Acknowledgments

We thank Professor Manuel Prieto for sharing with us his theoretical expression for the anisotropy decay due to energy migration in a tetramer.

## 8. Author Contributions

NK collected data and edited manuscript. AHAC analysed data and wrote the manuscript.

## References

[1] Lemmon, M. A. & Schlessinger, J. Cell signaling by receptor tyrosine kinases. Cell 141, 1117–1134, doi:10.1016/j.cell.2010.06.011 (2010).

[2] Normanno, N. et al. Epidermal growth factor receptor (EGFR) signaling in cancer. Gene 366, 2–16 (2006).

[3] Seshacharyulu, P. et al. Targeting the EGFR signaling pathway in cancer therapy. Expert Opinion on Therapeutic Targets 16, 15–31, doi:10.1517/14728222.2011.648617 (2012).

[4] Yarden, Y. and Schlessinger, J. Self-phosphorylation of epidermal growth factor receptor: evidence for a model of intermolecular allosteric activation. Biochemistry 26(5), 1434–42 (1987).

[5] Gadella, T.W. Jr and Jovin, T.M. Oligomerization of epidermal growth factor receptors on A431 cells studied by time-resolved fluorescence imaging microscopy. A stereochemical model for tyrosine kinase receptor activation. Journal of Cell Biology, 129(6), 1543–58 (1995).

[6] Clayton, A.H.A. et al. Ligand-induced dimer-tetramer transition during the activation of the cell surface epidermal growth factor receptor-A multidimensional microscopy analysis. Journal of Biological Chemistry, 280(34), 30392–9 (2005).

[7] Clayton, A.H., Orchard, S.G., Nice, E.C., Posner, R.G. and Burgess A.W. Predominance of activated EGFR higher-order oligomers on the cell surface. Growth Factors, 26(6), 316–24, (2008).

[8] Kozer, N., Barua, D., Orchard, S., Nice, E.C., Burgess, A.W., Hlavacek, W.S. and Clayton, A.H. Exploring higher-order EGFR oligomerisation and phosphorylation--a combined experimental and theoretical approach. Molecular Biosystems, 9(7), 1849–63 (2013).

[9] Kozer, N., Barua, D., Henderson, C., Nice, E.C., Burgess, A.W., Hlavacek, W.S. and Clayton, A.H. Recruitment of the adaptor protein Grb2 to EGFR tetramers. Biochemistry, 53(16), 2594–604 (2014).

[10] Needham, S. R. et al. EGFR oligomerization organizes kinase-active dimers into competent signalling platforms. Nature Communications, 7, 13307, doi:10.1038/ncomms13307 (2016).

[11] Huang, Y. et al. Molecular basis for multimerization in the activation of the epidermal growth factor receptor. eLife 5, doi:10.7554/eLife.14107 (2016).

[12] Garrett, T. P. et al. Crystal structure of a truncated epidermal growth factor receptor extracellular domain bound to transforming growth factor alpha. Cell 110, 763–773 (2002).

[13] Ogiso, H. et al. Crystal structure of the complex of human epidermal growth factor and receptor extracellular domains. Cell 110, 775–787 (2002).

[14] Cho, H.S and Leahy, D.J. Structure of the extracellular region of HER3 reveals an interdomain tether. Science 297, 1330–33 (2002).

[15] Bocharov, E.V, Bragin, P.E., Pavlov, K.V., Bocharova, O.V., Mineev, K.S., Polyansky, A.A., Volynsky, P.E., Efremov, R.G., Arseniev, A.S. The Conformation of the Epidermal Growth Factor Receptor Transmembrane Domain Dimer Dynamically Adapts to the Local Membrane Environment. Biochemistry, 56(12), 1697–1705 (2007).

[16] Mineev, K.S., Bocharov, E.V., Pustovalova, Y.E., Bocharova, O.V., Chupin, V.V., Arseniev, A.S. Spatial structure of the transmembrane domain heterodimer of ErbB1 and ErbB2 receptor tyrosine kinases. J. Mol. Biol., 400, 231–43 (2010).

[17] Thiel, K.W., Carpenter, G. Epidermal growth factor receptor juxtamembrane region regulates allosteric tyrosine kinase activation. PNAS 104, 19238–43 (2007).

[18] Jura, N., Endres, N.F., Engel, K., Deindl, S., Das, R., et al. Mechanism for activation of the EGF receptor catalytic domain by the juxtamembrane segment. Cell, 137, 1293–307 (2009).

[19] Stamos, J., Sliwkowski, M.X., Eigenbrot, C. Structure of the epidermal growth factor receptor kinase domain alone and in complex with a 4-anilinoquinazoline inhibitor. J. Biol. Chem., 277, 46265–72 (2002).

[20] Zhang, X., Gureasko, J., Shen, K., Cole, P.A., Kuriyan, J. An allosteric mechanism for activation of the kinase domain of epidermal growth factor receptor. Cell, 125, 1137–49 (2006).

[21] Lee, N.Y. and Koland, J.G. Conformational changes accompany phosphorylation of the epidermal growth factor receptor C-terminal domain. Protein Sci. 14(11), 2793–803 (2005).

[22] Lee, N.Y., Hazlett, T.L. and Koland, J.G. Structure and dynamics of the epidermal growth factor receptor C-terminal phosphorylation domain. Protein Sci. 15(5), 1142–52. (2006).

[23] Keppel, T.R., Sarpong, K., Murray, E.M., Monsey, J., Zhu, J., Bose, R. Biophysical Evidence for Intrinsic Disorder in the C-terminal Tails of the Epidermal Growth Factor Receptor (EGFR) and HER3 Receptor Tyrosine Kinases. J Biol Chem., 292(2), 597–610 (2017).

[24] Kozer, N., Rothacker, J., Burgess, A.W., Nice, E.C., Clayton, A.H. Conformational dynamics in a truncated epidermal growth factor receptor ectodomain. Biochemistry, 50(23), 5130–9 (2011).

[25] Rigby, A.C., Barber, K.R., Shaw, G.S. and Grant, C.W. Transmembrane region of the epidermal growth factor receptor: behavior and interactions via 2H NMR. Biochemistry, 35(38), 12591–601 (1996).

[26] Kaplan, M. et al. EGFR Dynamics Change during Activation in Native Membranes as Revealed by NMR. Cell 167, 1241-+, doi:10.1016/j.cell.2016.10.038 (2016).

[27] Stein, R.A., Hustedt, E.J., Staros, J.V., Beth, A.H. Rotational dynamics of the epidermal growth factor receptor. Biochemistry 41(6), 1957–64 (2002).

[28] Zidovetzki, R., Yarden, Y., Schlessinger, J., Jovin, T.M. Microaggregation of hormone-occupied epidermal growth factor receptors on plasma membrane preparations. EMBO J. 5(2), 247–50. (1986).

[29] Lu, C., Mi, L.Z., Schürpf, T., Walz, T., Springer, T.A. Mechanisms for kinase-mediated dimerization of the epidermal growth factor receptor. J Biol Chem. 287(45), 38244–53. (2012).

[30] Kozer, N., Kelly, MP., Orchard, S., Burgess, A.W., Scott, A.M. and Clayton, A.H. Differential and synergistic effects of epidermal growth factor receptor antibodies on unliganded ErbB dimers and oligomers. Biochemistry 50(18), 3581–90 (2011).

[31] Carter, R.E. and Sorkin, A. Endocytosis of functional epidermal growth factor receptor-green fluorescent protein chimera. J Biol Chem. 273(52), 35000–7 (1998).

[32] Paila, Y.D., Kombrabail, M., Krishnamoorthy, G., Chattopadhyay A. Oligomerization of the serotonin(1A) receptor in live cells: a time-resolved fluorescence anisotropy approach. J Phys Chem B. 115(39), 11439–47. doi:10.1021/jp201458h (2011).

[33] Kozer, N., Clayton, A.H.A. Analysis of complex anisotropy decays from single-frequency polarized-phasor ellipse plots. Methods Appl Fluoresc. 4(2), 024005. doi:10.1088/2050-6120/4/2/024005. (2016)

[34] Spencer, R.D. and Weber, G. Influence of Brownian rotations and energy transfer upon the measurements of fluorescence lifetime. J. Chem. Phys. 52, 1654 (1970).

[35] Jameson, D.M., Gratton, E. and Hall, R. D. The measurement and analysis of heterogeneous emissions by multifrequency phase and modulation fluorometry. Appl. Spectrosc. Rev. 20, 55–106 (1984).

[36] Lakowicz, J. R.and Prendergast, F. G. Quantitation of hindered rotations of diphenylhexatriene in lipid bilayers by differential polarized phase fluorometry. Science 200, 1399–401 (1978).

[37] Clayton, A.H.A. The polarized AB plot for the frequency-domain analysis and representation of fluorophore rotation and resonance energy homotransfer. J Microsc. 232(2) (2008).

[38] Bialik, C.N., Wolf, B., Rachofsky, E.L., Ross, J.B., Laws, W.R. Dynamics of biomolecules: assignment of local motions by fluorescence anisotropy decay. Biophys J. 75(5), 2564–73 (1998).

[39] Clayton, A.H., Hanley, Q.S., Arndt-Jovin, D.J., Subramaniam, V., Jovin, T.M. Dynamic fluorescence anisotropy imaging microscopy in the frequency domain (rFLIM). Biophys J. 83(3), 1631–49 (2002).

[40] Bader, A.N., Hofman, E.G., Voortman, J., en Henegouwen, P.M. and Gerritsen, H.C. Homo FRET imaging enables quantification of protein cluster sizes with subcellular resolution. Biophys J. 97(9), 2613–22. doi:10.1016/j.bpj.2009.07.059. (2009).

[41] Kovacs E, Das R, Wang Q, Collier TS, Cantor A, Huang Y, Wong K, Mirza A, Barros T, Grob P, Jura N, Bose R, Kuriyan J. Analysis of the Role of the C-Terminal Tail in the Regulation of the Epidermal Growth Factor Receptor. Mol Cell Biol. 35(17), 3083–102 (2015).

[42] Cadena, D.L., Chan, C.L., Gill, G.N. The intracellular tyrosine kinase domain of the epidermal growth factor receptor undergoes a conformational change upon autophosphorylation. J Biol Chem. 269(1), 260–5 (1994).

[43] Mi, L.Z., Lu, C., Li, Z., Nishida, N., Walz, T., Springer, T.A. Simultaneous visualization of the extracellular and cytoplasmic domains of the epidermal growth factor receptor. Nat Struct Mol Biol., 18(9), 984–9. (2011). doi:10.1038/nsmb.2092.

[44] Yang K. S., Ilagan M. X., Piwnica-Worms D., Pike L. J. Luciferase fragment complementation imaging of conformational changes in the EGF receptor. J. Biol. Chem. 284, 7474–7482 (2009).

[45] Weber, G. Theory of fluorescence depolarization by anisotropic Brownian rotations. Discontinuous distribution approach. J. Chem. Phys. 55, 2399–2411 (1971).

[46] Piston, D.W., Gratton E. Orientational exchange approach to fluorescence anisotropy decay. Biophys J. 56(6), 1083–91 (1989).

[47] Liang, S.I. et al. Phosphorylated EGFR dimers are not sufficient to activate Ras. Cell Rep. 22(10), 2593–2600 (2018).

[48] Hiroshima, M. et al. Transient Acceleration of Epidermal Growth Factor Receptor Dynamics Produces Higher-Order Signaling Clusters. J. Mol. Biol. 430 (9), 1386–1401 (2018).

[49] Zanetti-Domingues, L.C. et al. The architecture of EGFR's basal complexes reveals autoinhibition and activation mechanisms in dimers and oligomers. Nature Communications(in press).

